# A Novel Plasmid-Encoded Mobile Colistin Resistance Gene *mcr-13.1* Detected in *Escherichia coli* Isolated from Grassland

**DOI:** 10.64898/2026.07.21.739773

**Authors:** Marwa Alawi, Thuy Thi Do, Fiona Brennan, Catherine M. Burgess, Fiona Walsh

**Affiliations:** Maynooth University; Department of Agriculture, Food and the Marine; Teagasc Food Research Centre Ashtown; Teagasc - The Irish Agriculture and Food Development Authority

**Keywords:** Mobile colistin resistance, plasmids, grass, antimicrobial resistance

## Abstract

Plasmid-encoded mobile colistin resistance (*mcr*) genes have raised concern due to dissemination potential. While *mcr* variants are reported across One Health compartments, they remain unreported in grass. This study characterises a novel *mcr* variant (*mcr-13.1*), detected in *Escherichia coli* isolated from the grass phylosphere within an agricultural grassland. The two *mcr*-positive isolates were clonal copies isolated at timepoints eight weeks apart. They belonged to the serotype O17:H18 and were of the sequence type ST394. The *E. coli* were phenotypically susceptible to β-lactams, aminoglycosides, quinolones, sulphonamides, phenicols, tetracyclines, diaminopyrimidine and colistin (Minimum Inhibitory Concentration (MIC) = 0.5 µg/mL). The *mcr-13.1* gene was encoded on an IncFIB plasmid. This plasmid was transferable by conjugation but the colistin MIC of the *E. coli* J53 transconjugant did not change (0.5 µg/mL). Further, cloned pUC19::*mcr-13.1* did not alter the colistin MIC for *E. coli* DH5α (0.25 µg/mL). The translated amino acid sequence showed highest homology (82 %) to MCR-10.2 and MCR-10.4. Our findings identify grass as a previously unrecognised reservoir for *E. coli* carrying mobile *mcr* genes, reports the identification of the novel *mcr-13.1* variant from this niche and demonstrates the importance of genomic screening in identifying *mcr* genes that would otherwise remain undetected.

## INTRODUCTION

Antimicrobial resistance (AMR) has emerged as a global problem that is described as one of the top ten public health issues facing the human population ^1^. Extended-spectrum β-lactamase (ESBL) and carbapenemase-producing Enterobacterales, multidrug-resistant *Pseudomonas aeruginosa* and carbapenem-resistant *Acinetobacter baumannii* have emerged as pathogens of highest priority due to limited treatment options ^2^. This has led to the resurgence in the use of polymyxin antimicrobials which were once discontinued due to reported nephrotoxicity. Polymyxins are cationic cyclic polypeptide antimicrobials which comprise of polymyxin B and polymyxin E (colistin), amongst others ^3^. Colistin is a last-resort polymyxin antimicrobial often reserved for the treatment of multidrug-resistant Gram-negative bacterial infections. However, colistin-resistance has increased in recent years.

Colistin resistance in Enterobacterales has been primarily due to chromosomal mutations e.g. two-component systems PhoPQ and PmrAB ^4^. Recently, mobile colistin resistance (*mcr*) has emerged and is a cause of concern, as *mcr* genes are often on conjugative or mobilisable plasmids, which can move between and across species and environments ^5^. The *mcr* genes encode the MCR family of phosphoethanolamine transferases (PEts). The predominant theory regarding the mechanism of action is that the PEts catalyse the addition of phosphoethanolamine onto lipid A in the outer membrane of Gram-negative bacteria. This results in a reduction of the negative charge, which prevents the efficient binding of colistin and thus leading to resistance or reduced susceptibility ^6^.

The first report of a plasmid-encoded *mcr* gene (*mcr-1*) was identified in *E. coli* and *Klebsiella pneumoniae* from animal and human sources in China ^7^. Since the discovery of *mcr-1*, ten homologs (*mcr-1 – mcr-10*) and their variants have been reported in bacteria isolated from across humans, animals and environmental sectors ^8,9^, with an additional two (*mcr-11* and *mcr-12*) deposited in the NCBI Reference Gene Catalogue but not yet described in the literature. In Ireland, *mcr-1, mcr-8* and *mcr-9* have been reported in agricultural, clinical, and environmental samples associated with *K. pneumoniae, E. coli* and *Enterobacter sp.* ^10,11^. This study describes both the first global report of a novel *mcr* variant (*mcr-13.1*) and the detection of *mcr* in *E. coli* isolated from grass.

## RESULTS

### Sampling & Antimicrobial Susceptibility Testing

Two *E. coli* isolates were recovered from agricultural grass in Ireland, 8-weeks apart. Both isolates were susceptible to all antimicrobials tested (ampicillin, cefotaxime, ceftazidime, imipenem, kanamycin, amikacin, gentamicin, ciprofloxacin, sulphonamide, chloramphenicol, tetracycline, trimethoprim and colistin).

### Genomic Analysis

The two *E. coli* strains belonged to the sequence type ST394 and serogroup O17:H18. *E. coli* Ec_018 was isolated at week 2 of sampling and *E. coli* Ec_325 was isolated eight weeks later at week 10. Ec_018 was isolated from grass within a plot that had been treated with fresh manure, whereas Ec_325 was isolated from untreated grass. Average nucleotide identity assessment confirmed a 99.99 % similarity between the two isolates. This indicates persistence of the same clone over at least an 8-week period. Genome screening initially revealed the presence of an *mcr-9.1*-like gene sharing a 75.94 % nucleotide identity with the reference *mcr-9.1* from *Salmonella enterica* (accession: [NZ_NAAN01000000](https://www.ncbi.nlm.nih.gov/nuccore/NZ_NAAN00000000.1)). The translated amino acid sequence of the *mcr-13.1* gene shared 76.95 % amino acid identity with this reference. However, further investigation revealed that our translated MCR-13.1 protein shared 81.78 % sequence similarity with the reference MCR-10.2 and MCR-10.4 (Table 1). Both reference variants originated in *Enterobacter sp.* from human sources. MCR-10.4 was associated with *Enterobacter ludwigii* from a human throat and MCR-10.2 was linked to *Enterobacter kobei* from human secretion.

**Table 1:**
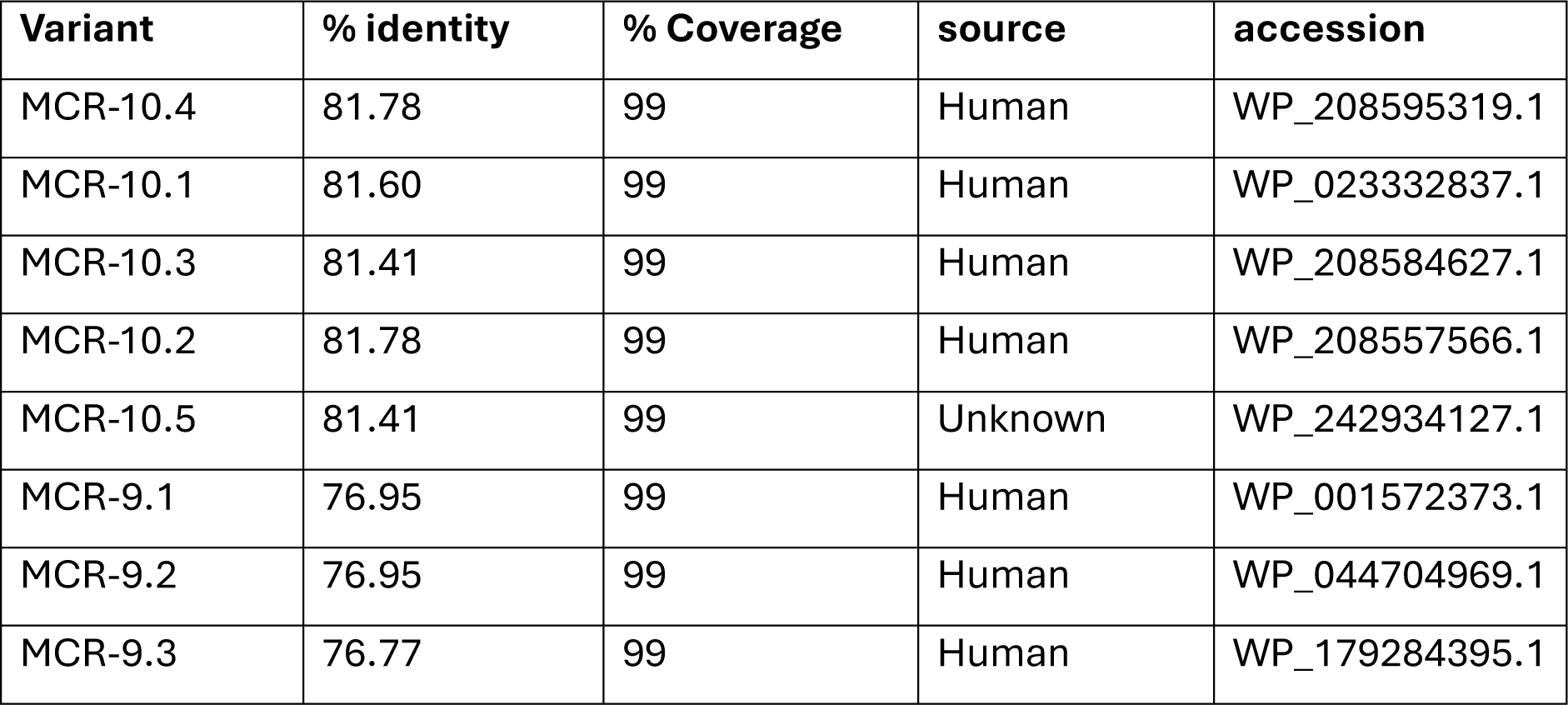
MCR alleles sharing the closest amino acid sequence identity to our MCR-13.1 variant.

| Variant | % identity | % Coverage | source | accession |
| --- | --- | --- | --- | --- |
| MCR-10.4 | 81.78 | 99 | Human | WP_208595319.1 |
| MCR-10.1 | 81.60 | 99 | Human | WP_023332837.1 |
| MCR-10.3 | 81.41 | 99 | Human | WP_208584627.1 |
| MCR-10.2 | 81.78 | 99 | Human | WP_208557566.1 |
| MCR-10.5 | 81.41 | 99 | Unknown | WP_242934127.1 |
| MCR-9.1 | 76.95 | 99 | Human | WP_001572373.1 |
| MCR-9.2 | 76.95 | 99 | Human | WP_044704969.1 |
| MCR-9.3 | 76.77 | 99 | Human | WP_179284395.1 |

Generally, phylogenetic analysis grouped MCR subvariants into distinct clades (e.g. MCR-1 family, MCR-2 etc.). Our MCR-13.1 variant (Figure 1, orange) shared ancestry with the MCR-9 and MCR-10 clades but branched out from the MCR-10 family to form its own clade (Figure ).

**Figure 1:**
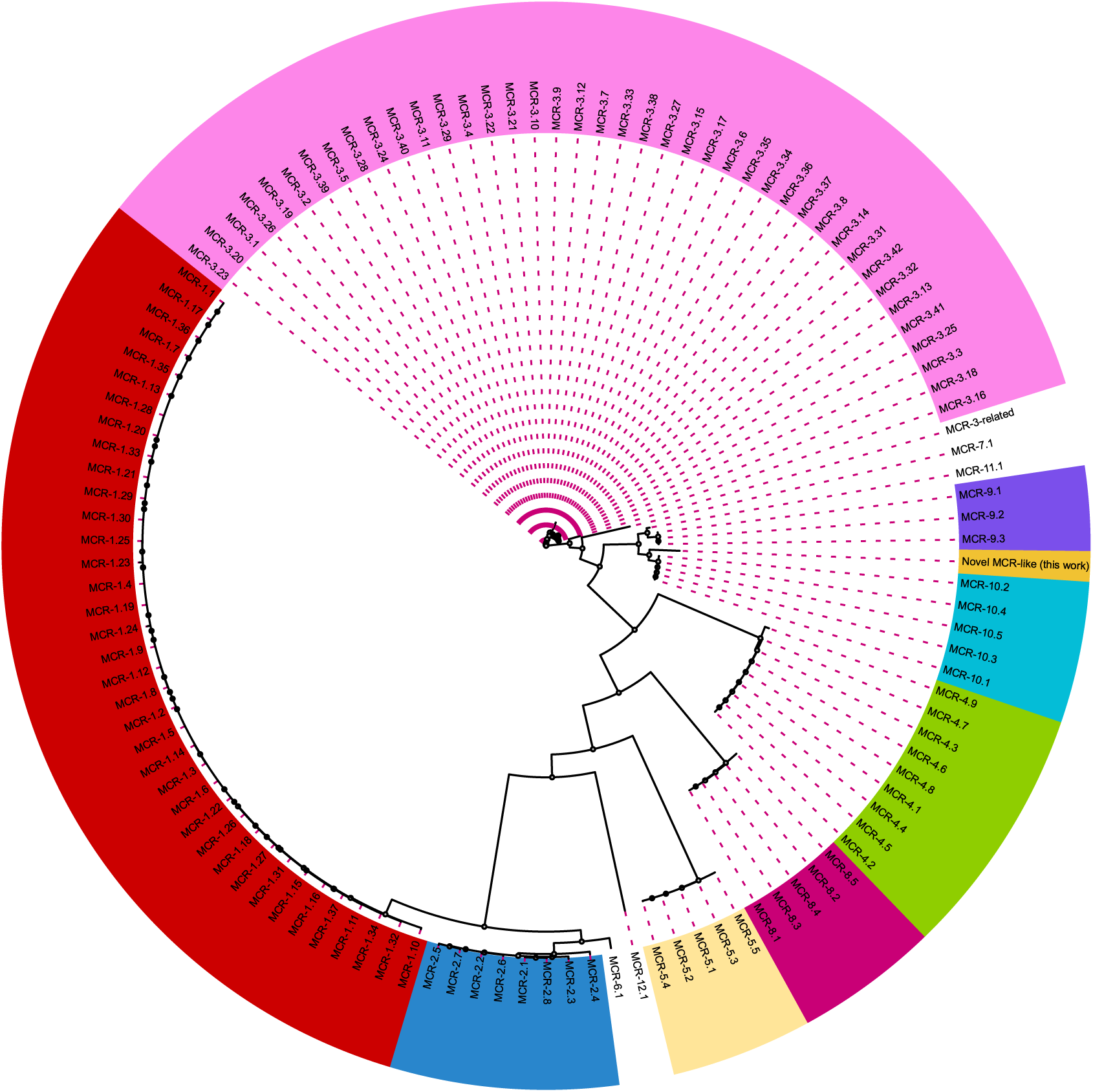
Phylogenetic placement of the novel MCR variant based on amino acid sequences. Maximum-likelihood phylogenetic tree inferred from aligned amino acid sequences of all MCR protein variants deposited in NCBI. The translated MCR-13.1 amino acid sequence identified in this study is highlighted in orange to demonstrate its evolutionary relationship to established lineages. The MCR-13.1 sequence shares ancestry with MCR-10 variants. It shares 82 % sequence similarity with the MCR-10.2 and MCR-10.4 variants, suggesting evolutionary divergence from the MCR-10 family.

The relatively low amino acid sequence similarity and the departure from known MCR clades suggested that our variant may be encoding a divergent MCR phosphoethanolamine transferase. No additional antimicrobial resistance genes were detected. However, genome screening suggested the presence of a plasmid replicon sharing 83.07 % identity with IncFIB(pB171)_1_pB171 in both isolates. Short-read assemblies predicted the *mcr* gene was located on IncFIB-type plasmid contigs.

### Identification of Homologous Amino Acid Sequences

Sequences matching the MCR-13.1 were identified through a BLASTp search. These shared 99.26 % identity with MCR-13.1 and were reportedly isolated from *E. coli* from a wildlife centre (accession: HAY0261769.1) and *Cronobacter sakazakii* (accessions: ELY4545964.1) from a bathroom floor.

### Colistin MIC & *mcr-13.1* mobility

Both isolates were susceptible to colistin (MIC = 0.5 µg/mL) indicating a divergence in genotype and phenotype. However, the *mcr-13.1*-encoding plasmid was successfully transferred into recipient *E. coli* J53, confirming its conjugative nature. The colistin MICs of the transconjugants showed no increase compared to the recipient *E. coli* J53 (MIC = 0.5 µg/mL), indicating that the *mcr* gene did not alter the MIC.

### Plasmid Localisation & Genetic Context

Long-read sequencing using MinION ONT resolved a 116,256 bp IncFIB plasmid carrying the *mcr-13.1* gene. Two virulence factors were encoded on this plasmid: a serine protease precursor autotransporter (*pic)* and the F4 fimbriae (f*aeC*). The plasmid encoded a MOBF relaxase and shared a MASH neighbour distance of 0.113 with the nearest MASH neighbour (accession: CP025891). The distance suggests that our plasmid is potentially novel or not well represented in the databases.

The *mcr-13.1* gene was flanked by a small hypothetical gene (141 bp) upstream and an IS3 transposase (300 bp) downstream (Figure B). An ISRoR2 transposase (942 bp) was located upstream of the hypothetical gene, while an IS256 family transposase (219 bp) was present downstream of IS3. The tyrosine recombinase gene *xerC* was also present upstream of the ISRoR2. The presence of a tyrosine recombinase gene and multiple insertion sequences flanking the *mcr* gene suggests a mobilizable genetic environment.

**Figure 2:**
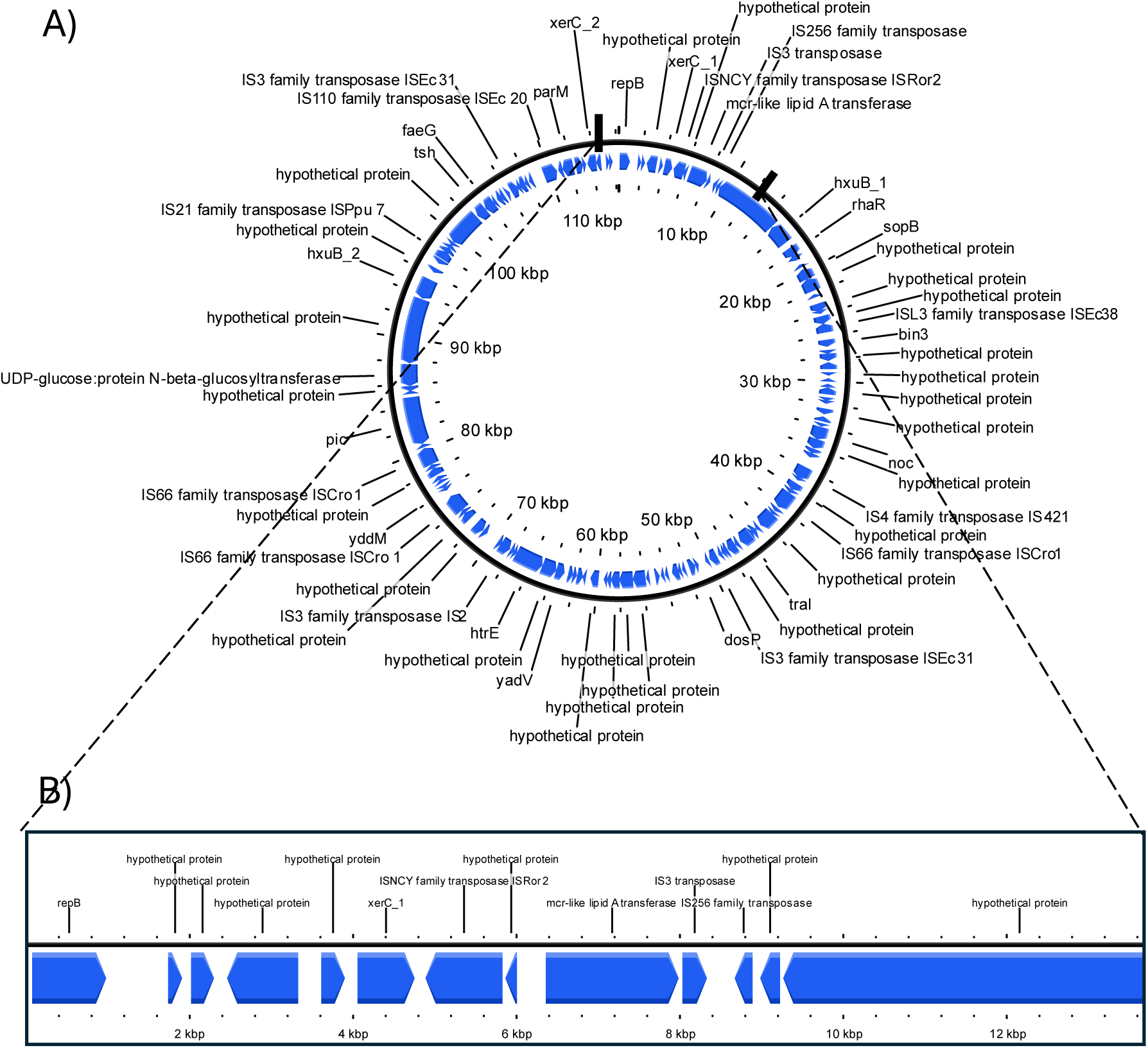
Plasmid map displaying coding sequences (CDS) annotated by Prokka. A) Overview of all annotated CDS, B) Zoomed-in view of CDS flanking the *mcr* gene. The CDS are represented by blue arrows indicating transcriptional direction. In addition to insertion sequences, other CDS of note are the *repB* and *tra* genes, indicators of a conjugative plasmid. A list of all genes is available in Supplementary Table 1.

### Inducibility of MCR expression

We screened for the two-component QseBC system that is often associated with *mcr-S* induction ^12^. However, this was absent from our genomes. Therefore, the *mcr-13.1* gene was cloned into the high-copy vector pUC19 with an upstream ribosomal binding site (RBS) to facilitate translation. The cloned vectors produced identical colistin MICs to the empty pUC19 vector (0.25 ug/mL). No variation in MIC was observed upon IPTG induction.

### Amino Acid Structural Comparison

Predicted structures generated using the AlphaFold server (Figure ) show that the overall fold of our MCR protein is highly similar to that of MCR-10.2 and MCR-10.4. All three proteins adopt a similar 3D structure, with conserved α-helices and β-sheets secondary structures.

**Figure 3:**
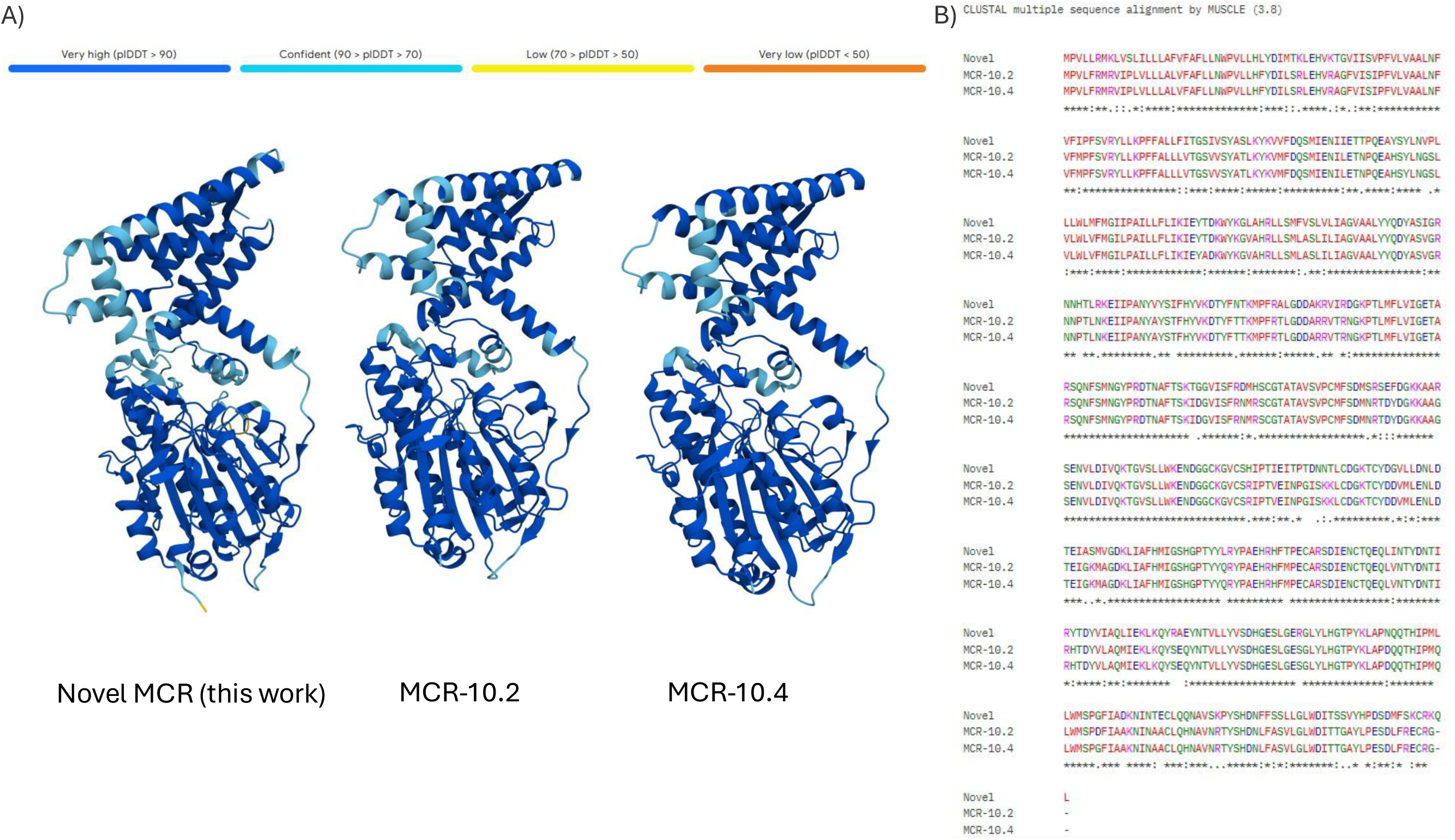
Structural and sequence comparison of the MCR protein and its closest homologs. A) Predicted 3D structures of the MCR-13.1 described in this study and its closest homologs (MCR-10.2 and MCR-10.4). B) Multiple Sequence Alignment of the MCR protein and its homologs generated by MUSCLE. Conserved residues are indicated by (*), strong and weak similarities are indicated by (: and .) respectively. The absence of a symbol is indicative of non-conserved residues.

This corroborated the multiple sequence alignment which revealed a high degree of conservation between the three variants, with highly conserved regions indicated by the “*” symbol. However, the MCR we describe in this study shows a total of 13 non-conserved positions. These substitutions are distributed throughout the protein sequence suggesting localised variability and moderate evolutionary divergence from MCR-10.2 and MCR-10.4.

## DISCUSSION

In this study, we describe a novel *mcr-13.1* gene located on a conjugative IncFIB plasmid of two *E. coli* isolates recovered from grass in Ireland. The isolates were highly clonal, belonging to ST394 and were of the O17:H18 group. *E. coli* ST394 has been isolated primarily from animal sources such as bovines and goats ^13–15^. The recovery of an almost identical lineage of *E. coli* ST394 over an eight-week interval suggests ecological stability within the grassland environment. No farm animals had been grazed on the field for seven months prior to sampling. The recovery of this *E. coli* clone from both manured and non-manured grass confirms that the isolate was not introduced via manure application. *E. coli* ST394 from bovine and goats has been previously linked to the *mcr-1.1* variant. This was chromosomally-encoded in one *E. coli* and carried on an IncHI2 plasmid in another ^14,15^.

Although the isolates described in this study were phenotypically susceptible to colistin, genomic analysis revealed the presence of a putative phosphoethanolamine transferase sharing 82 % amino acid similarity with MCR-10.2 and MCR-10.4 originally described in *Enterobacter sp.* ^16^. Generally, the MCR subvariants cluster closely within the same clade due to minor substitutions (point mutations) that distinguishes each subvariants within a family ^17^. The phosphoethanolamine transferase identified in this study shows a level of similarity to MCR-10.2 and MCR-10.4; however, its placement as a distinct branch within the MCR-10 clade suggests that it may represent a novel allele diverging from this MCR family. This divergence appears to be driven by multiple non-conserved amino acid substitutions which may be contributing to subtle differences in structure and expression. These non-conserved substitutions are not clustered within a specific functional domain but rather are dispersed throughout the translated amino acid sequence, indicating a pattern of gradual sequence divergence. Two strains deposited in the NCBI, an *E. coli* from a wildlife centre and *C. sakazakii* from a bathroom floor contained highly similar versions of our MCR-13.1 variant, indicating that this allele may already be circulating, highlighting the importance of characterising allelic diversity in surveillance studies. Notably, our work represents the first genomic description of *mcr-13.1* and its function in the context of colistin-resistance.

The presence of this novel phosphoethanolamine transferase on a conjugative plasmid represents a One Health threat due to its potential to spread via horizontal gene transfer. Our work demonstrated the *mcr-13.1* gene is transferable via a conjugative IncFIB plasmid (116,256 bp), alongside virulence genes, including *pic* and *faeC*. IncFIB plasmids are frequently identified in *Enterobacterales* from all dimensions of One Health and are often linked with virulence genes, as well as antimicrobial resistance ^18,19^. Plasmids encoding *mcr* genes are commonly associated with multiple resistance genes, including ESBL and carbapenemase genes ^20,21^ facilitating co-selection under various conditions. This is widely reported across diverse plasmid incompatibility groups and bacterial hosts, highlighting the role of *mcr-*encoding plasmids in the dissemination of multidrug-resistance. In contrast, the plasmid characterised in our work is atypical as it did not harbour additional resistance genes; this was reflected in the susceptible phenotypes against all tested antimicrobial classes. The absence of additional resistance genes may be an evolutionary response to reducing the metabolic burden associated with *mcr* genes ^22,23^ allowing for enhanced stability in the absence of selective pressures.

Insertion sequences IS3 and ISRoR2 transposases flank the *mcr-13.1* gene and may have contributed to its mobility ^24^. IS3 transposases have previously been linked to other *mcr* variants, particularly *mcr-3* ^25–28^ but have not yet been directly linked with *mcr-9* or *-10* variants. ISRoR2 transposases have not previously been reported in association with *mcr* genes. However, *C. sakazakii* (accession no.: ABPDRR010000037.1), which encodes for a phosphoethanolamine transferase sharing 99.26 % amino acid identity with our transferase, encodes a putative transposase that shares 99.77 % identity with our ISRoR2 transposase suggesting a potentially related genetic context.

Site-specific recombination and transposition may have also contributed to the mobilisation and integration of our *mcr-13.1* gene into the IncFIB plasmid. XerC-type tyrosine recombinase is often reported immediately upstream of *mcr-10* ^16,29–31^. In our plasmid, the ISNCY family transposase ISRoR2 interjects between a XerC tyrosine recombinase and the *mcr-13.1* variant. However, the presence of XerC in close proximity to the *mcr* suggests a potential role in stabilising the ORF in our IncFIB plasmid ^32^.

The *mcr-13.1* gene did not confer colistin resistance when present on the IncFIB plasmid nor when cloned into pUC19. In its original plasmid context, the gene neither conferred resistance in the grass *E. coli* nor increased the MIC in the *E. coli* J53 transconjugant. This reflects *mcr-9* genes which are not associated with changes in colistin tolerance or resistance ^33,34^. The QseBC two-component system has been linked to inducible *mcr-9* expression at subinhibitory concentrations of colistin ^12^. However, this system was absent from our *E. coli* chromosomes and plasmids. On the other hand, the *mcr-10* variants confer colistin-resistance or reduced susceptibility when cloned into expression vectors ^29,30^, which was not the case for our variant. Similarly, induction of the pUC19 *lac* promoter did not alter the colistin MIC, indicating that increased expression of our variant did not affect the resistance phenotype. The absence of induction-dependent increase in MIC is consistent with previous observations for *mcr-10* (Wang *et al.*, 2020). It remains unclear why this gene has been maintained within the IncFIB plasmid.

In conclusion, we report the identification of a novel mobile *mcr-13.1* variant and have identified the first *mcr* in grass-derived *E. coli*. Functional characterisation did not reveal a colistin-resistance phenotype, suggesting that the *mcr-13.1* may be non-functional or not expressed under the conditions tested. This study highlights the importance of both the functional characterisation and genomic screening for the identification of novel variants within the *mcr* gene families and proposes that grass is a potential reservoir for the spread and dissemination of *mcr* genes in *E. coli*.

## METHODS

### Sampling and Processing

Two *E. coli* isolates were recovered from grass samples collected in Ireland during the summer in 2019. The grassland site was in the Teagasc research facility in Johnstown Castle, Co. Wexford, Ireland (52.294117°N, −6.50102°W). Grass samples were collected as previously described ^35^ as part of a slurry land spreading experiment. Briefly, presumptive *E. coli* were selected on Eosin Methylene Blue (EMB) agar according to green metallic sheen morphology. Sampling took place every two weeks starting before slurry application, then two weeks later at week 0, week 2, week 4, week 6 and week 10. The two *E. coli* described in this study were selected at week 2 and week 10 of sampling (i.e. 8 weeks apart) chosen based on genomic analysis described below. *E. coli* Ec_018 was isolated from fresh pig slurry -treated grass at week 2 of sampling, while *E. coli* Ec_325 was later isolated from untreated grass at week 10 of sampling.

### Antimicrobial Susceptibility Testing

The Kirby-Bauer disk testing method using the Clinical & Laboratory Standards Institute (CLSI) guidelines was performed ^36^. Both isolates were tested for susceptibility to ampicillin (10 μg), cefotaxime (30 μg), ceftazidime (30 μg), imipenem (10 μg), gentamicin (10 μg), amikacin (30 μg), kanamycin (30 μg), tetracycline (30 μg), sulfonamide (300 μg), trimethoprim (5 μg), chloramphenicol (30 μg), and ciprofloxacin (5 μg). Quality control strains *E. coli* ACTC 25922 and *Pseudomonas aeruginosa* ATCC 27853 were used as references. Colistin resistance was assessed using microbroth dilutions following CLSI guidelines. Quality control strains *P. aeruginosa* ATCC 27853 and *E. coli* NCTC 13846 (*mcr-1*) were negative and positive controls, respectively.

### DNA Extraction & Whole Genome Sequence

Microbial DNA was extracted from both *E. coli* strains using the Macherey Nagel Nucleospin Microbial DNA kit according to manufacturer instructions. Whole genome sequencing was performed using short-read Illumina sequencing by Novogene.

The UseGalaxy server (https://usegalaxy.eu/) was used for data processing. Raw files were trimmed using TrimGalore! v0.6.10 (https://github.com/FelixKrueger/TrimGalore) and assembled with Unicycler v0.5.1 (https://github.com/rrwick/Unicycler). The assembled contigs were quality assessed using Quast v5.3.0 (https://github.com/ablab/quast). Separately, on the KBase platform (https://www.kbase.us/), CheckM v1.0.18 (https://github.com/Ecogenomics/CheckM) was used to define high quality genome completeness (≥90 % completeness, <5 % contamination). Species identity was confirmed using the “identify species” function on PubMLST https://pubmlst.org/. Average Nucleotide Identification (ANI) was used to determine the genetic relatedness of the two *E. coli* isolates.

Genomes were annotated using Prokka v1.14.6 (https://github.com/tseemann/prokka) using the default settings. The sequence types and serogroups were identified using MLST (https://github.com/tseemann/mlst) and EcOH (https://github.com/katholt/srst2), respectively. Screening was performed using ABRicate for ARGs and plasmid replicon types using the comprehensive antimicrobial resistance database (CARD), PointFinder, ResFinder and PlasmidFinder. For all databases, the minimum coverage was at default setting of 80 %. The identity was reduced to 70 % to identify potential distance homologs and to account for potential database bias towards clinical samples.

To identify whether the *mcr* variant was present on the chromosome or a plasmid, both genomes were further processed using PlasFlow to separate chromosomes from plasmids. The predicted plasmid sequences were re-annotated using ABRicate (CARD, ResFinder) with reduced identity score of 70 %.

### In Silico Identification of Homologous Amino Acid Sequences

The translated amino acid sequence of the *mcr-13.1* gene was compared against publicly available sequences using BLASTp to determine if this gene had previously been deposited in public databases. Sequence matches were evaluated based on percentage identity, query coverage and corresponding species.

### Nucleotide sequence accession numbers

The genome assemblies of Ec_018 and Ec_352 have been deposited into GenBank under the accession no. GCF_056783495.1 and GCA_056783455.1, respectively.

The assemblies can be accessed at: https://www.ncbi.nlm.nih.gov/assembly/GCA_056783495.1

https://www.ncbi.nlm.nih.gov/assembly/GCA_056783455.1

### Long read plasmid sequencing

Plasmid DNA was extracted using the Macherey Nagel-Nucleospin plasmid purification kit using the low-copy number protocol as per manufacturer’s instructions. The extracted plasmid DNA was sequenced using Oxford Nanopore Technology (ONT) MinION with the native barcoding kit (SQK-NBD114). The raw fast5 files were quality controlled, base called and demultiplexed using Dorado v7.9.8. On the UseGalaxy platform, sequencing adapters were removed with Porechop v0.2.4 without fastq outputs (https://github.com/rrwick/porechop) followed by an additional QC filter step using Filtlong v0.3.1 (https://github.com/rrwick/Filtlong) to exclude short read sequences (contigs <1000 bp). Plasmids were then assembled using Unicycler v0.5.1 with long reads only. Reads were visualised using Bandage v2022.09. As with WGS, the plasmids were annotated using Prokka and screened using ABRicate with reduced identity percentage (70 %). In addition, MOB-recon at default settings was used for typing and reconstructing plasmids. The genetic context of the *mcr* open reading frame (ORF) was predicted using Prokka-generated annotations. Where possible, ORFs reported as “hypothetical” were annotated using BLAST similarity prediction.

### Evolutionary Contextualisation

To visualise the evolutionary ancestors of our identified protein, all MCR amino acid sequences were downloaded from NCBI. A maximum-likelihood phylogenetic tree was produced using FastTree (https://github.com/morgannprice/fasttree). The phylogenetic tree was then visualised on iTOL v7 (https://itol.embl.de/). The AlphaFold Server (https://alphafoldserver.com/) was used to predict the structure of the MCR protein and its closest two homologs based on amino acid sequences. Multiple sequence alignment was powered by MUSCLE (https://www.ebi.ac.uk/jdispatcher/msa) with ClustalW output on all three proteins to identify conserved and non-conserved regions.

### Mobility of *mcr-13.1* Gene

To confirm if *mcr-13.1* was mobilisable, conjugation assays were performed using the original environmental *E. coli* as the plasmid donor with the recipient strain *E. coli* J53 (azide-resistant). Presumptive transconjugants were selected on Mueller-Hinton agar supplemented with 150 µg/mL sodium azide and colistin at 0, 1 and 2 µg/mL. The environmental *E. coli* was confirmed to be susceptible to sodium azide at this concentration. Presumptive transconjugants grew at 1 µg/mL. As colistin does not diffuse well in agar, a random selection of presumptive transconjugant colonies underwent confirmatory *mcr* PCR using forward 5’-TGACCAAGCTCGAACACGT-3’ and reverse primer 5’-AATGACACCGCCAGTTTTGC-3’ to produce a ∼700 bp amplicon.

### Transconjugant Susceptibility Testing

Colistin microbroth dilution susceptibility tests were performed on the *E. coli* J53 transconjugants confirmed to be *mcr*-positive, *E. coli* J53 wild type and the controls using CLSI guidelines as described previously.

### Cloning of *mcr-13.1* Gene

The entire coding region (1626 bp) of the *mcr-13.1* gene was amplified from the environmental *E. coli*using forward 5’-ATA<u>AAGCTT</u>AGGAGGTTTTAATATGCCCGTACTATTAAGGATGA -3’ and reverse 5’-ATA<u>GGATCC</u>TTAGCGACACTTGCTGAACATATC-3’ primers. HindIII and BamHI restriction sites (underlined) were incorporated in the forward and reverse primers, respectively. A canonical ribosomal-binding site (AGGAGG) and a spacer (TTTTAAT) was added 6 bp upstream of the forward primer to facilitate translation. The insert was cloned into the pUC19 vector using double digestion by BamHI-HF and HindIII-HF, followed by ligation at 16 ℃ overnight with T4 ligase. The cloned vector was subsequently electroporated into *E. coli* DH5α. Transformants were selected using blue-white screening on LB agar supplemented with X-gal (200 µg/mL) and isopropyl β-D-1-thiogalactopyranoside (IPTG, 1mM). Screening was performed in both the presence and absence of subinhibitory concentrations of colistin at 0.25 µg/mL. Resulting white colonies were screened by PCR to confirm successful transformation, as described previously. Colistin microbroth dilution susceptibility tests were repeated in the presence and absence of IPTG (1mM) for the DH5α *mcr-13.1-*transformants, wild type DH5α, DH5α-pUC19, and control strains *P. aeruginosa* ATCC 27853 and *E. coli* NCTC 13846 (*mcr-1*).

### Verification of pUC1G::*mcr-13.1* Constructs by Sequencing

The cloned vectors (n = 3) were extracted from DH5α pUC19::*mcr-13.1* using the Macherey-Nagel Plasmid Purification kit following the high-copy number protocol as per manufacturer’s protocols. Vectors were sequenced using ONT by Novogene, UK.

## ACKNOWLEDGEMENT

We thank the staff who contributed and managed the field trial in the Teagasc field centre in Wexford. The authors would like to thank Dr. Meghana Srinivas for advice and support regarding the sequencing components of this work.

## AUTHOR CONTRIBUTIONS

**M. Alawi:** Conceptualization, methodology, software, data curation, writing - original draft preparation, visualization, investigation. **T. T. Do:** methodology, sample acquisition. **F. Brennan:** Supervision, Project Administration, Resources. **C Burgess**: Project Administration, Resources, Supervision. **F. Walsh:** Conceptualization, writing-review and editing, supervision, funding acquisition.

## DATA AVAILABILITY

The assembled sequences for Ec_018 and Ec_325 have been deposited in the NCBI GenBank and are publicly available under the accession numbers GCA_056783495.1 (Ec_018) and GCA_056783455.1 (Ec_325).

The assemblies can be accessed at: https://www.ncbi.nlm.nih.gov/assembly/GCA_056783495.1

https://www.ncbi.nlm.nih.gov/assembly/GCA_056783455.1

## REFERENCES

1. Naghavi, M. et al. Global burden of bacterial antimicrobial resistance 1990–2021: a systematic analysis with forecasts to 2050. The Lancet 404, 1199–1226 (2024).

2. Word Health Organisation. WHO Bacterial Priority Pathogens List 2024: Bacterial Pathogens of Public Health Importance, to Guide Research, Development, and Strategies to Prevent and Control Antimicrobial Resistance. (World Health Organization, Geneva, 2024).

3. Evans, M. E., Feola, D. J. & Rapp, R. P. Polymyxin B Sulfate and Colistin: Old Antibiotics for Emerging Multiresistant Gram-Negative Bacteria. Ann. Pharmacother. 33, 960–967 (1999).

4. Olaitan, A. O., Morand, S. & Rolain, J.-M. Mechanisms of polymyxin resistance: acquired and intrinsic resistance in bacteria. Front. Microbiol. 5, 643 (2014).

5. Anyanwu, M. U., Jaja, I. F. & Nwobi, O. C. Occurrence and Characteristics of Mobile Colistin Resistance (mcr) Gene-Containing Isolates from the Environment: A Review. Int. J. Environ. Res. Public. Health 17, 1028 (2020).

6. Hinchliffe, P. et al. Insights into the Mechanistic Basis of Plasmid-Mediated Colistin Resistance from Crystal Structures of the Catalytic Domain of MCR-1. Sci. Rep. 7, 39392 (2017).

7. Liu, Y.-Y. et al. Emergence of plasmid-mediated colistin resistance mechanism MCR-1 in animals and human beings in China: a microbiological and molecular biological study. Lancet Infect. Dis. 16, 161–168 (2016).

8. Gharaibeh, M. H. & Shatnawi, S. Q. An overview of colistin resistance, mobilized colistin resistance genes dissemination, global responses, and the alternatives to colistin: A review. Vet. World 12, 1735–1746 (2019).

9. Hussein, N. H., AL-Kadmy, I. M. S., Taha, B. M. & Hussein, J. D. Mobilized colistin resistance (mcr) genes from 1 to 10: a comprehensive review. Mol. Biol. Rep. 48, 2897–2907 (2021).

10. Cahill, N. et al. First reported detection of the mobile colistin resistance genes, *mcr*-8 and *mcr*-9, in the Irish environment. Sci. Total Environ. 876, 162649 (2023).

11. Smyth, C., Leigh, R. J., Do, T. T. & Walsh, F. Communities of plasmids as strategies for antimicrobial resistance gene survival in wastewater treatment plant effluent. Npj Antimicrob. Resist. 3, 78 (2025).

12. Kieffer, N. et al. mcr-9, an Inducible Gene Encoding an Acquired Phosphoethanolamine Transferase in Escherichia coli, and Its Origin. Antimicrob. Agents Chemother. 63, 10.1128/aac.00965-19 (2019).

13. Hassen, B., et al. *mcr*-1 encoding colistin resistance in CTX-M-1/CTX-M-15-producing *Escherichia coli* isolates of bovine and caprine origins in Tunisia. First report of CTX-M-15-ST394/D *E. coli* from goats. Comp. Immunol. Microbiol. Infect. Dis. 67, 101366 (2019).

14. Naha, S. et al. Carriage and within-host diversity of mcr-1.1-harbouring Escherichia coli from pregnant mothers: inter- and intra-mother transmission dynamics of mcr-1.1. Emerg. Microbes Infect. 12, 2278899 (2023).

15. Haenni, M. et al. No genetic link between *E. coli* isolates carrying *mcr-1* in bovines and humans in France. J. Glob. Antimicrob. Resist. 41, 111–116 (2025).

16. Liao, W. et al. High prevalence of colistin resistance and *mcr-9/10* genes in *Enterobacter* spp. in a tertiary hospital over a decade. Int. J. Antimicrob. Agents 59, 106573 (2022).

17. Partridge, S. R. et al. Proposal for assignment of allele numbers for mobile colistin resistance (mcr) genes. J. Antimicrob. Chemother. 73, 2625–2630 (2018).

18. Rozwandowicz, M. et al. Plasmids carrying antimicrobial resistance genes in Enterobacteriaceae. J. Antimicrob. Chemother. 73, 1121–1137 (2018).

19. Smyth, C., Leigh, R. J., Delaney, S., Murphy, R. A. & Walsh, F. Shooting hoops: globetrotting plasmids spreading more than just antimicrobial resistance genes across One Health. Microb. Genomics 8, mgen000858 (2022).

20. Lakshmanan, D., Ramasamy, D., Subramanyam, V. & Saravanan, S. K. Mobile colistin resistance (mcr) genes and recent developments in colistin resistance detection. Lett. Appl. Microbiol. 76, ovad102 (2023).

21. Zhang, Q. Bacteria carrying mobile colistin resistance genes and their control measures, an updated review. Arch. Microbiol. 206, 462 (2024).

22. Liu, Y.-Y. et al. Metabolic Perturbations Caused by the Over-Expression of mcr-1 in Escherichia coli. Front. Microbiol. 11, 588658 (2020).

23. Liang, L. et al. Fitness costs of mobilised colistin resistance gene 3 (mcr-3): systematic review, epidemiological study, and functional analysis. eBioMedicine 120, 105923 (2025).

24. Vandecraen, J., Chandler, M., Aertsen, A. & Van Houdt, R. The impact of insertion sequences on bacterial genome plasticity and adaptability. Crit. Rev. Microbiol. 43, 709–730 (2017).

25. Eichhorn, I. et al. Identification of novel variants of the colistin resistance gene mcr-3 in Aeromonas spp. from the national resistance monitoring programme GERM-Vet and from diagnostic submissions. J. Antimicrob. Chemother. 73, 1217–1221 (2018).

26. Wang, X. et al. Presence of an mcr-3 Variant in Aeromonas caviae, Proteus mirabilis, and Escherichia coli from One Domestic Duck. Antimicrob. Agents Chemother. 62, e02106–17 (2018).

27. Xiang, R. et al. Colocation of the Polymyxin Resistance Gene mcr-1 and a Variant of mcr-3 on a Plasmid in an Escherichia coli Isolate from a Chicken Farm. Antimicrob. Agents Chemother. 62, e00501–18 (2018).

28. Trongjit, S. & Chuanchuen, R. Whole genome sequencing and characteristics of Escherichia coli with co-existence of ESBL and mcr genes from pigs. PLoS ONE 16, e0260011 (2021).

29. Wang, C. et al. Identification of novel mobile colistin resistance gene mcr-10. Emerg. Microbes Infect. 9, 508–516 (2020).

30. Xu, T. et al. Identification of mcr-10 carried by self-transmissible plasmids and chromosome in Enterobacter roggenkampii strains isolated from hospital sewage water. Environ. Pollut. 268, 115706 (2021).

31. Guan, J. et al. First Report of the Colistin Resistance Gene mcr-10.1 Carried by IncpA1763-KPC Plasmid pSL12517-mcr10.1 in Enterobacter cloacae in Sierra Leone. Microbiol. Spectr. 10, e01127–22 (2022).

32. Castillo, F., Benmohamed, A. & Szatmari, G. Xer Site Specific Recombination: Double and Single Recombinase Systems. Front. Microbiol. 8, 453 (2017).

33. Tyson, G. H. et al. The mcr-9 Gene of Salmonella and Escherichia coli Is Not Associated with Colistin Resistance in the United States. Antimicrob. Agents Chemother. 64, 10.1128/aac.00573-20 (2020).

34. Lemlem, M. et al. Phenotypic and genotypic characterization of colistin-resistant Escherichia Coli with mcr-4, mcr-5, mcr-6, and mcr-9 genes from broiler chicken and farm environment. BMC Microbiol. 23, 392 (2023).

35. Do, T. T. et al. Comparison of soil and grass microbiomes and resistomes reveals grass as a greater antimicrobial resistance reservoir than soil. Sci. Total Environ. 857, 159179 (2023).

36. CLSI. CLSI M100-ED36:2026 Performance Standards for Antimicrobial Susceptibility Testing. https://em100.edaptivedocs.net/GetDoc.aspx?doc=CLSI%20M100%20ED36:2026&scope=user (2026).

