## Supplemental Table 1 for "A Novel Plasmid-Encoded Mobile Colistin Resistance Gene *mcr-13.1* Detected in *Escherichia coli* Isolated from Grassland"

| locus_tag | ftype | length_bp | gene | EC_numbe | COG | product | Author Comments |
|---|---|---|---|---|---|---|---|
| JAIEELLP_00001 | CDS | 906 | repB | | | RepFIB replication protein A | |
| JAIEELLP_00002 | CDS | 171 | | | | hypothetical protein | |
| JAIEELLP_00003 | CDS | 279 | | | | hypothetical protein | |
| JAIEELLP_00004 | CDS | 870 | | | | hypothetical protein | |
| JAIEELLP_00005 | CDS | 288 | | | | hypothetical protein | |
| JAIEELLP_00006 | CDS | 699 | xerC_1 | | | Tyrosine recombinase XerC | |
| JAIEELLP_00007 | CDS | 942 | | | | ISNCY family transposase ISRor2 | |
| JAIEELLP_00008 | CDS | 141 | | | | hypothetical protein | |
| JAIEELLP_00009 | CDS | 1626 | mcr-9.1 | | | phosphoethanolamine--lipid A transferase MCR-9.1 | now known as "*mcr-13.1* " |
| JAIEELLP_00010 | CDS | 300 | | | | hypothetical protein | "IS3 transposase" based on BLAST search |
| JAIEELLP_00011 | CDS | 219 | | | | hypothetical protein | "IS256 family transposase" based on BLAST sea |
| JAIEELLP_00012 | CDS | 243 | | | | hypothetical protein | |
| JAIEELLP_00013 | CDS | 5784 | | | | hypothetical protein | |
| JAIEELLP_00014 | CDS | 1800 | hxuB_1 | | COG2831 | Heme/hemopexin transporter protein HuxB | |
| JAIEELLP_00015 | CDS | 837 | rhaR | | | HTH-type transcriptional activator RhaR | |
| JAIEELLP_00016 | CDS | 105 | | | | hypothetical protein | |
| JAIEELLP_00017 | CDS | 291 | | | | hypothetical protein | |
| JAIEELLP_00018 | CDS | 291 | | | | hypothetical protein | |
| JAIEELLP_00019 | CDS | 960 | sopB | | | Protein SopB | |
| JAIEELLP_00020 | CDS | 1167 | | | | hypothetical protein | |
| JAIEELLP_00021 | CDS | 303 | | | | hypothetical protein | |
| JAIEELLP_00022 | CDS | 708 | | | | hypothetical protein | |
| JAIEELLP_00023 | CDS | 153 | repA | | | Replication protein RepA | |
| JAIEELLP_00024 | CDS | 708 | | | | hypothetical protein | |
| JAIEELLP_00025 | CDS | 795 | | | | ISL3 family transposase ISEc38 | |

| | | | | | | |
|---|---|---|---|---|---|---|
| JAIEELLP_00026 | CDS | 621 | bin3 | | | Putative transposon Tn552 DNA-invertase bin3 |
| JAIEELLP_00027 | CDS | 336 | | | | hypothetical protein |
| JAIEELLP_00028 | CDS | 648 | | | | hypothetical protein |
| JAIEELLP_00029 | CDS | 609 | | | | hypothetical protein |
| JAIEELLP_00030 | CDS | 201 | | | | hypothetical protein |
| JAIEELLP_00031 | CDS | 135 | | | | hypothetical protein |
| JAIEELLP_00032 | CDS | 228 | | | | hypothetical protein |
| JAIEELLP_00033 | CDS | 627 | | | | hypothetical protein |
| JAIEELLP_00034 | CDS | 201 | | | | hypothetical protein |
| JAIEELLP_00035 | CDS | 309 | | | | hypothetical protein |
| JAIEELLP_00036 | CDS | 507 | | | | hypothetical protein |
| JAIEELLP_00037 | CDS | 441 | | | | hypothetical protein |
| JAIEELLP_00038 | CDS | 240 | | | | hypothetical protein |
| JAIEELLP_00039 | CDS | 1134 | noc | | | Nucleoid occlusion protein |
| JAIEELLP_00040 | CDS | 846 | | | | hypothetical protein |
| JAIEELLP_00041 | CDS | 432 | psiB | | | Protein PsiB |
| JAIEELLP_00042 | CDS | 399 | | | | hypothetical protein |
| JAIEELLP_00043 | CDS | 1113 | | | COG3385 | IS4 family transposase IS421 |
| JAIEELLP_00044 | CDS | 141 | | | | hypothetical protein |
| JAIEELLP_00045 | CDS | 678 | | | | hypothetical protein |
| JAIEELLP_00046 | CDS | 348 | | | | hypothetical protein |
| JAIEELLP_00047 | CDS | 1572 | | | | IS66 family transposase ISCro1 |
| JAIEELLP_00048 | CDS | 342 | | | | hypothetical protein |
| JAIEELLP_00049 | CDS | 282 | traQ | | | Protein TraQ |
| JAIEELLP_00050 | CDS | 1431 | | | | hypothetical protein |
| JAIEELLP_00051 | CDS | 279 | traD | | | Coupling protein TraD |
| JAIEELLP_00052 | CDS | 399 | vapC_1 | 3.1.-.- | COG1487 | tRNA(fMet)-specific endonuclease VapC |
| JAIEELLP_00053 | CDS | 228 | vapB_1 | | COG4456 | Antitoxin VapB |

| | | | | | | |
|---|---|---|---|---|---|---|
| JAIEELLP_00054 | CDS | 1059 | traI | | | Multifunctional conjugation protein TraI |
| JAIEELLP_00055 | CDS | 333 | | | | hypothetical protein |
| JAIEELLP_00056 | CDS | 741 | | | | hypothetical protein |
| JAIEELLP_00057 | CDS | 372 | | | | hypothetical protein |
| JAIEELLP_00058 | CDS | 216 | finO | | | Fertility inhibition protein |
| JAIEELLP_00059 | CDS | 312 | | | | hypothetical protein |
| JAIEELLP_00060 | CDS | 678 | | | | IS3 family transposase ISEc31 |
| JAIEELLP_00061 | CDS | 213 | | | | hypothetical protein |
| JAIEELLP_00062 | CDS | 228 | dosP | 3.1.4.52 | COG2199 | Oxygen sensor protein DosP |
| JAIEELLP_00063 | CDS | 552 | pdeG_1 | 3.1.4.52 | | putative cyclic di-GMP phosphodiesterase PdeG |
| JAIEELLP_00064 | CDS | 213 | | | | hypothetical protein |
| JAIEELLP_00065 | CDS | 156 | | | | hypothetical protein |
| JAIEELLP_00066 | CDS | 591 | | | | hypothetical protein |
| JAIEELLP_00067 | CDS | 270 | | | | hypothetical protein |
| JAIEELLP_00068 | CDS | 126 | | | | hypothetical protein |
| JAIEELLP_00069 | CDS | 387 | | | | hypothetical protein |
| JAIEELLP_00070 | CDS | 159 | | | | hypothetical protein |
| JAIEELLP_00071 | CDS | 1116 | | | | hypothetical protein |
| JAIEELLP_00072 | CDS | 1209 | | | | hypothetical protein |
| JAIEELLP_00073 | CDS | 762 | | | | hypothetical protein |
| JAIEELLP_00074 | CDS | 369 | | | | hypothetical protein |
| JAIEELLP_00075 | CDS | 252 | ftsE | | COG2884 | Cell division ATP-binding protein FtsE |
| JAIEELLP_00076 | CDS | 720 | lnt | 2.3.1.269 | | Apolipoprotein N-acyltransferase |
| JAIEELLP_00077 | CDS | 216 | | | | ISNCY family transposase ISLad2 |
| JAIEELLP_00078 | CDS | 513 | | | | hypothetical protein |
| JAIEELLP_00079 | CDS | 351 | | | | hypothetical protein |

| | | | | | | |
|---|---|---|---|---|---|---|
| JAIEELLP_00080 | CDS | 435 | | | | hypothetical protein |
| JAIEELLP_00081 | CDS | 765 | yadV | | | putative fimbrial chaperone YadV |
| JAIEELLP_00082 | CDS | 1284 | | | | hypothetical protein |
| JAIEELLP_00083 | CDS | 2520 | htrE | | COG3188 | Outer membrane usher protein HtrE |
| JAIEELLP_00084 | CDS | 273 | | | | hypothetical protein |
| JAIEELLP_00085 | CDS | 327 | | | | hypothetical protein |
| JAIEELLP_00086 | CDS | 906 | | | | IS3 family transposase IS2 |
| JAIEELLP_00087 | CDS | 366 | | | | hypothetical protein |
| JAIEELLP_00088 | CDS | 396 | | | | hypothetical protein |
| JAIEELLP_00089 | CDS | 834 | | | | hypothetical protein |
| JAIEELLP_00090 | CDS | 678 | | | | hypothetical protein |
| JAIEELLP_00091 | CDS | 348 | | | | hypothetical protein |
| JAIEELLP_00092 | CDS | 1572 | | | | IS66 family transposase ISCro1 |
| JAIEELLP_00093 | CDS | 282 | yddM | | COG3093 | putative HTH-type transcriptional regulator YddM |
| JAIEELLP_00094 | CDS | 282 | higB | 3.1.-.- | | Endoribonuclease HigB |
| JAIEELLP_00095 | CDS | 309 | | | | hypothetical protein |
| JAIEELLP_00096 | CDS | 351 | | | | ISNCY family transposase ISLad2 |
| JAIEELLP_00097 | CDS | 678 | | | | hypothetical protein |
| JAIEELLP_00098 | CDS | 348 | | | | hypothetical protein |
| JAIEELLP_00099 | CDS | 1572 | | | | IS66 family transposase ISCro1 |
| JAIEELLP_00100 | CDS | 111 | | | | hypothetical protein |
| JAIEELLP_00101 | CDS | 4242 | pic | 3.4.21.- | | Serine protease pic autotransporter |
| JAIEELLP_00102 | CDS | 237 | | | | hypothetical protein |
| JAIEELLP_00103 | CDS | 441 | | | | hypothetical protein |
| JAIEELLP_00104 | CDS | 1911 | | 2.4.1.- | | UDP-glucose:protein N-beta-glucosyltransferase |

| | | | | | | |
|---|---|---|---|---|---|---|
| JAIEELLP_00105 | CDS | 5784 | | | | hypothetical protein |
| JAIEELLP_00106 | CDS | 1800 | hxuB_2 | | COG2831 | Heme/hemopexin transporter protein HuxB |
| JAIEELLP_00107 | CDS | 411 | | | | hypothetical protein |
| JAIEELLP_00108 | CDS | 414 | | | | hypothetical protein |
| JAIEELLP_00109 | CDS | 768 | | | | IS21 family transposase ISPpu7 |
| JAIEELLP_00110 | CDS | 450 | | | | hypothetical protein |
| JAIEELLP_00111 | CDS | 345 | | | | hypothetical protein |
| JAIEELLP_00112 | CDS | 174 | | | | hypothetical protein |
| JAIEELLP_00113 | CDS | 3030 | | | | hypothetical protein |
| JAIEELLP_00114 | CDS | 915 | tsh | 3.4.21.- | | Temperature-sensitive hemagglutinin tsh autotransporter |
| JAIEELLP_00115 | CDS | 885 | faeG | | | K88 fimbrial protein AD |
| JAIEELLP_00116 | CDS | 267 | | | | hypothetical protein |
| JAIEELLP_00117 | CDS | 411 | pdeL | 3.1.4.52 | COG2200 | Cyclic di-GMP phosphodiesterase PdeL |
| JAIEELLP_00118 | CDS | 378 | pdeG_2 | 3.1.4.52 | | putative cyclic di-GMP phosphodiesterase PdeG |
| JAIEELLP_00119 | CDS | 312 | | | | hypothetical protein |
| JAIEELLP_00120 | CDS | 678 | | | | IS3 family transposase ISEc31 |
| JAIEELLP_00121 | CDS | 174 | | | | hypothetical protein |
| JAIEELLP_00122 | CDS | 543 | | | | hypothetical protein |
| JAIEELLP_00123 | CDS | 300 | papB | | | Major pilu subunit operon regulatory protein PapB |
| JAIEELLP_00124 | CDS | 348 | | | | hypothetical protein |
| JAIEELLP_00125 | CDS | 1197 | | | | IS110 family transposase ISEc20 |
| JAIEELLP_00126 | CDS | 393 | | | | hypothetical protein |
| JAIEELLP_00127 | CDS | 972 | parM | | | Plasmid segregation protein ParM |

| | | | | | | |
|---|---|---|---|---|---|---|
| JAIEELLP_00128 | CDS | 645 | srp54 | | | Signal recognition particle 54 kDa protein |
| JAIEELLP_00129 | CDS | 276 | | | | hypothetical protein |
| JAIEELLP_00130 | CDS | 783 | xerC_2 | | | Tyrosine recombinase XerC |
| JAIEELLP_00131 | CDS | 420 | | | | hypothetical protein |
| JAIEELLP_00132 | CDS | 231 | vapB_2 | | COG4456 | Antitoxin VapB |
| JAIEELLP_00133 | CDS | 312 | vapC_2 | 3.1.-.- | COG1487 | tRNA(fMet)-specific endonuclease VapC |
| JAIEELLP_00134 | CDS | 270 | | | | hypothetical protein |
| JAIEELLP_00135 | CDS | 426 | | | | hypothetical protein |

arch
